# SHOOT: phylogenetic gene search and ortholog inference

**DOI:** 10.1101/2021.09.01.458564

**Authors:** D.M. Emms, S. Kelly

**Author notes:** ***Corresponding Author***, Name: Steven Kelly, Name: David Emms.

## Abstract

Determining the evolutionary relationships between genes is fundamental to comparative biological research. Here we present the phylogenetic search, SHOOT. SHOOT searches a user query sequence against a database of phylogenetic trees and returns a tree with the query sequence correctly placed within it. We show that SHOOT performs this analysis with comparable speed to a BLAST search. We demonstrate that SHOOT phylogenetic placements are as accurate as conventional tree inference and it can identify orthologs with high accuracy. In summary, SHOOT is a fast and accurate tool for phylogenetic analyses of novel query sequences. It is available online at www.shoot.bio.

## Background

Resolving the phylogenetic relationships between biological sequences provides a framework for inferring sequence function, and a basis for understanding the diversity and evolution of life on Earth. The entry point to such phylogenetic analyses is provided by algorithms that either align or identify regions of local similarity between pairs of biological sequences. The first implementations of such algorithms utilised global alignments to provide a basis to score similarity between sequences [1]. Later, faster local alignment methods were developed [2], followed by the FASTA heuristic database search [3] and culminating with the development of the BLAST algorithm and statistical methods for homology testing [4] in the 1990s. Since then, BLAST and other local alignment methods [5–7] have provided a critical foundation of biological science research and form the entry point to the majority of biological sequence analyses.

One feature of the problem that is under-utilised in BLAST and related local alignment search tools is the transitive nature of homology. Because local alignment searching methods do not store the relationships between sequences, a search of a query gene against a large database will involve carrying out many needless pairwise local alignments against numerous closely related homologs. An alternative approach would be to infer the relationships between all database sequences ahead of time using phylogenetic inference methods. These phylogenetic relationships can then be stored as part of the database, facilitating the use of lighter-weight search approaches or sparse reference databases with relationships already computed. Existing methods that take these kind of approaches include TreeFam for genes within the Metazoa [8] and TreeGrafter for annotating protein sequences using annotated phylogenetic trees [9].

Although local similarity searches such as BLAST are the primary entry point to the sequence analysis, a frequent end-goal of such analyses is to identify orthologs of the query sequence in other species. The use of phylogenetic methods is the canonical method for assessing gene relationships. Phylogenetic methods for estimating sequence similarity are more accurate than using local pairwise alignments, and critically they provide contextual information about the place of the query gene within its gene family. This includes the identification of orthologs, paralogs, and gene gain and loss within each clade in of the resultant phylogenetic tree. Although the similarity scores returned by local alignment methods can be used to approximate phylogenetic trees [10], they are not accurate and can be limited by only having alignments against a single query gene rather than alignments between sequences already in the database [11]. Moreover, even when all pairwise similarity scores are calculated the accuracy of phylogenetic trees inferred from these scores is limited [10]

Here we present SHOOT, a software tool for rapidly searching a phylogenetically partitioned and structured database of biological sequences. There are a number of advantages to taking a phylogenetic approach to sequence searching. We show that by grouping homologous genes in the database, a gene can then be rapidly assigned to its homology group, irrespective of the number of homologous genes. Further, false negatives are unlikely since complete homology groups can be identified securely ahead of time. This helps avoid the reduced sensitivity that results from local sequence similarity database search algorithm heuristics used to determine which sequences to consider aligning [6]. Phylogenetic inference methods can then be used to rapidly and accurately assign the gene to its correct position within the otherwise pre-computed gene tree for its homology group [12]. This avoids the need to evaluate gene-relatedness using e-values, which are a measure of the certainty that a pair of genes are homologous, rather than a direct evaluation of the phylogenetic relationship between genes [13]. In summary, SHOOT efficiently and accurately places query sequences directly into phylogenetic trees. In this way the phylogenetic history of the query sequence and its orthologs can be immediately visualised, interpreted, and retrieved. SHOOT is provided for use at www.shoot.bio.

## Results

### Pre-computed databases of phylogenetic trees allow ultra-fast phylogenetic orthology analysis of novel gene sequences

The conventional procedure for sequence orthology analysis is to first assemble a group of gene sequences which share similarity and then perform phylogenetic tree inference on this group to infer the relationships between those genes. The SHOOT algorithm was designed to make such a phylogenetic analysis feasible as a real-time search using a two-stage approach. The first stage comprises the ahead-of-time construction of a SHOOT phylogenetic database and the second stage implements the SHOOT search for a query sequence (Figure 1). The database preparation phase includes multiple automated steps including homology group inference, multiple sequence alignment, phylogenetic tree inference, and homology group profiling (see Methods). Thus, prior to database searching the phylogenetic relationships between all genes in the database are already established. Subsequent SHOOT searches exploit the fact that the alignments and trees have already been computed to enable the use of accurate phylogenetic methods for placement of query genes within pre-computed gene trees with little extra computation required.

**Figure 1.**
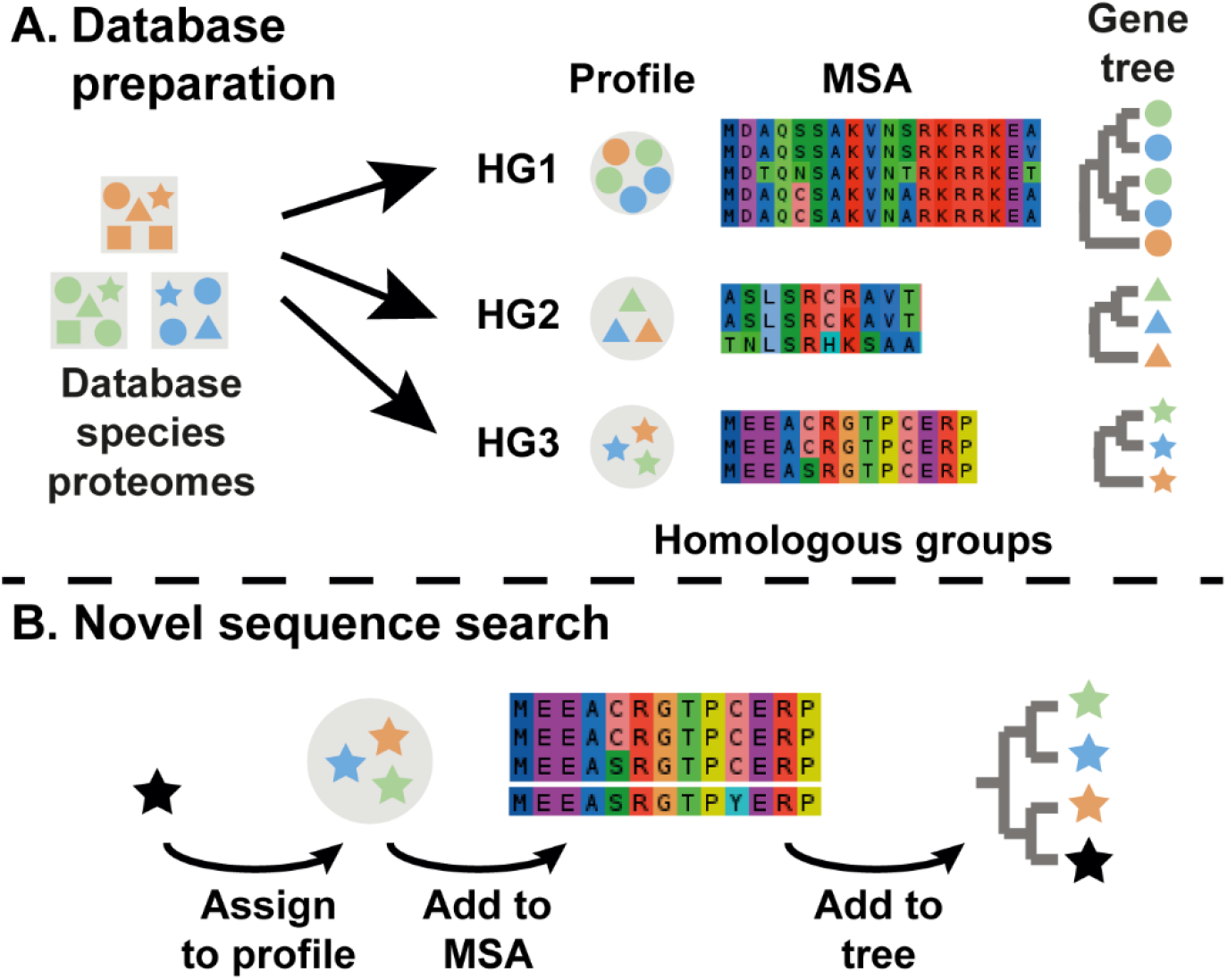
The workflow for the two separate stages of SHOOT: **A**) The database preparation stage. **B**) The sequence search stage. MSA, multiple sequence alignment. HG, homologous group. Individual shapes represent individual protein sequences.

The median time for a complete a SHOOT search of a database containing 984,137 protein sequences from 78 species was 5.5 seconds using 16 cores of an Intel Xeon E5-2683 CPU for (Figure 2A). This compared with 1.19 seconds for a conventional BLAST search of the same sequence set (Figure 2A). However, unlike BLAST (or similar) sequence search methods, the output of a SHOOT search is not an ordered list of similar sequences but is instead a maximum likelihood phylogenetic tree with bootstrap support values inferred from a multiple sequence alignment with the query gene embedded within it. SHOOT also computes the orthologs of the query gene using phylogenetic methods.

**Figure 2.**
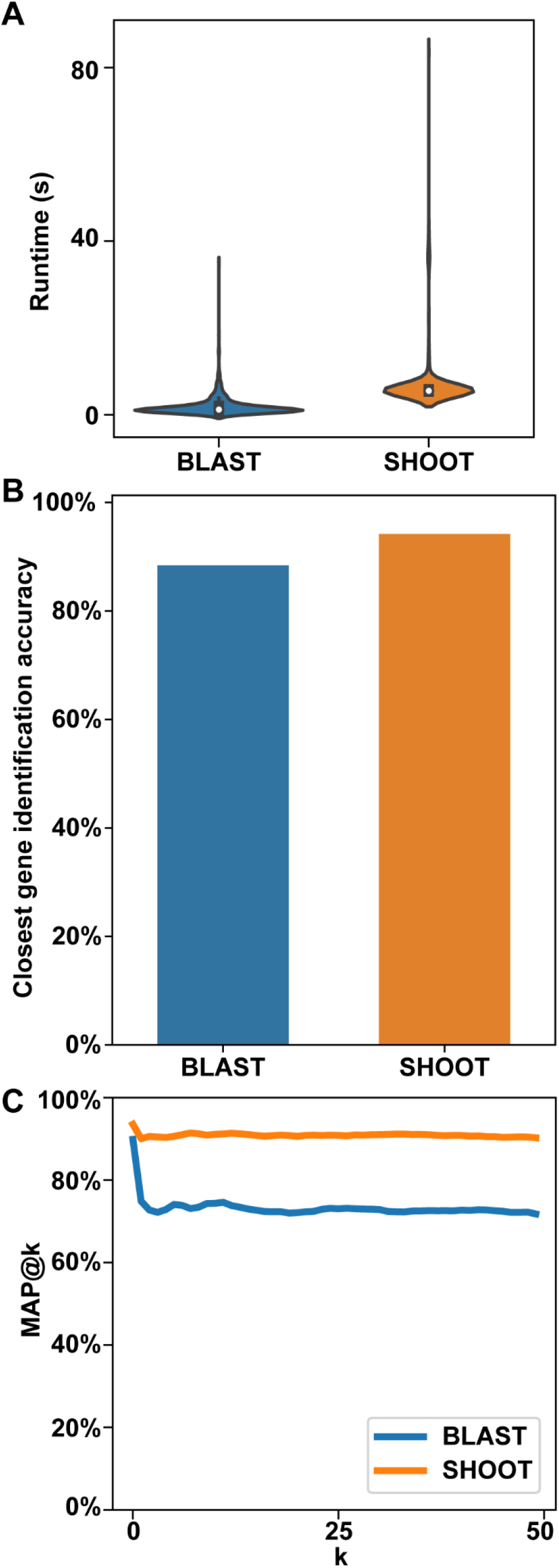
Runtime and closest homologs identification accuracy for SHOOT and BLAST. A) Violin plot of runtimes for 1000 searches of randomly sampled sequences against the same database of 984,137 protein sequences from 78 species. B) Accuracy at identifying the closest related database gene to a randomly selected query sequence. C) Mean Average Precision at k (MAP@k).

### SHOOT is more accurate than BLAST in identifying the closest related gene sequence

A leave-one-out analysis was conducted to test SHOOT’s ability to find the most closely related gene sequence in a given database. Here a set of 1000 test cases was randomly sampled from the UniProt Reference Proteomes database. Each test case consisted of a pair of genes sister to each other with at least 95% bootstrap support in a maximum likelihood gene tree. One member of the test pair was arbitrarily designated the “query sequence” and the other gene was designated “the expected closest gene” i.e. the gene that should be identified by a search method as the most similar gene in the database. To provide a comparison, BLAST [11] was also tested on the same dataset. The set of query genes were searched against the database and each method was scored on whether or not the closest/best scoring gene in each search result was “the expected closest gene”. The tests showed that SHOOT identified “the expected closest gene” as the most closely related gene in 94.2% of cases (Figure 2A). For comparison, BLAST correctly identified the “the expected closest gene” as the most similar gene sequence in 88.4% of cases. To put this in context, there is a 1 in 9 chance that the top hit returned by BLAST is not the most closely related sequence in the database while there is a 1 in 17 chance that the same is true for SHOOT. Thus, SHOOT is better able to identify the closest related gene to a given query gene in a given database and can be used as an alternative to BLAST for this purpose.

### SHOOT gives evolutionary context of a query gene’s position within its gene family

Although for many users knowledge of the closest related gene as described above may be sufficient, in many instances there will be more than one gene that is equally closely related to the query gene in a given species. Thus, to generalise the “best hit” analysis above for larger gene sets the “Mean Average Precision at k” score [14] was calculated, to quantify the precision at which the k closest homologs identified by SHOOT or BLAST correspond to the k expected closest homologs in maximum likelihood gene trees. This analysis was conducted for values of k between 1 (equivalent to the “best hit” analysis above) and 50 (Figure 2B). As k increased, MAP@k for BLAST fell to 71.8%. i.e. there was a 71.8% agreement between the closest homologs identified using BLAST and those identified using phylogenetic methods. In contrast, the use of phylogenetic methods in the database construction stage of SHOOT coupled with the accurate placement of genes within the database trees (Figure 2A), resulted in MAP@50 for SHOOT of 90.3%. Thus, both the list of most closely related genes and their rank order of relationship to the query gene is substantially more accurate for SHOOT than for BLAST.

### SHOOT has high accuracy in identifying orthologs of the query gene

A frequent goal of sequence similarity searches is to identify orthologs of the query gene in other species. As stated above, local similarity search tools such as BLAST do not do this. Instead, they return a list of genes that should be subject to multiple sequence alignment and phylogenetic inference in order to infer the orthology relationships between genes. The phylogenetic tree returned by SHOOT provides the evolutionary relationships between genes inferred from multiple sequence alignment and maximum likelihood tree inference allowing orthologs and paralogs to be identified. SHOOT also automatically identifies orthologs and colours the genes in the tree according to whether they are orthologs or paralogs (Supplementary Figure 1), as identified using the species overlap method [15, 16], which has been shown to be an accurate method for automated orthology inference [17]. The tree viewer also supports a zoom functionality to view a progressively larger or smaller clade of genes around the query gene. An image of the tree can be downloaded, the tree can also be exported in Newick format, and the FASTA file of protein sequences in the tree can be downloaded to support further downstream analyses.

To evaluate the accuracy of ortholog inference 6 species were chosen at increasing time since divergence from human. These query species comprised Mouse, Chicken, Zebrafish, the Tunicate *Ciona intestinalis*, fruit fly, and the yeast *Saccharomyces cerevisiae* (Figure 3A). Orthologs between these species and Human were determined from OrthoFinder on the 2020 Quest for Orthologs benchmark dataset [13, 17]. For each query species 100 query genes were selected, creating a test set of 600 genes in total. For these 600 genes SHOOT was evaluated on its accuracy in identifying the orthologs in human. For comparison BLAST best hit (BH) and reciprocal best hit (RBH) were likewise evaluated (Figure 3B). SHOOT was between 11% (Mouse) and 47% (*S. cerevisiae*) more accurate than either method using BLAST and the difference was greatest for more diverged species (Figure 3B). The greatest difference between SHOOT and BLAST was in the percentage of orthologs that were recovered (Recall, Figure 3C). For all species, the ortholog recall for SHOOT was >79%. Whereas the ortholog recall for BLAST RBH was for 37% for *S*. cerevisiae, the most distant species from human in the analysis (Figure 3C). The precision of SHOOT orthologs was intermediate between BLAST RBH and BH (Figure 3D). Thus, SHOOT ortholog assignments are more accurate than performing a “top hit” or “reciprocal best BLAST hit” analysis for identification of orthologs.

**Figure 3.**
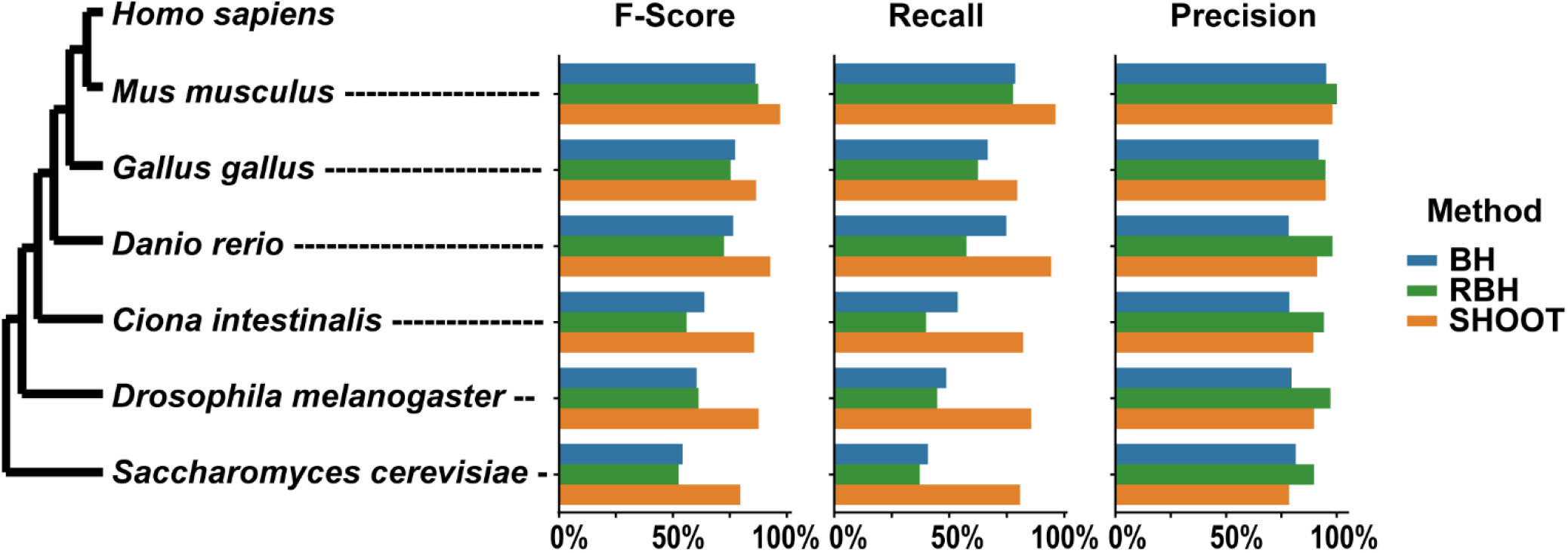
F-score, precision and recall at identifying orthologs in Homo sapiens for 100 query genes in each of Mus musculus, Gallus gallus, Danio rerio, Ciona intestinalis, Drosophila melanogaster and Saccharomyces cerevisiae for BLAST best hit (BH), BLAST reciprocal best hit (RBH) and SHOOT.

### Curated databases place the gene in the context of model species and key events in the gene’s evolution

The initial release of SHOOT includes phylogenetic databases for Metazoa, Fungi, Plants, Bacteria & Archaea, and also the UniProt Quest for Orthologs (QfO) reference proteomes, which cover all domains of cellular life (Supplementary Tables 1–5). To maximise the utility of the gene trees to a wide range of researchers, the species within the databases have been chosen to contain model species, species of economic or scientific importance, and species selected because of their key location within the evolutionary history covered by the database. Each database also contains multiple outgroup species to allow robust rooting of the set of gene trees. As an example, Supplementary Figure 2 shows the phylogeny for the metazoan database, highlighting the taxonomic groups of the included species. Although a number of databases are provided on the SHOOT webserver, the SHOOT command line tool has been designed so that databases can be compiled from any species set.

## Discussion and Conclusions

SHOOT is a phylogenetic search engine for analysis of biological sequences. It has been designed to take a user-provided query sequence and return a phylogenetic analysis of that sequence using a database of reference organisms. We show that SHOOT can perform this search and analysis with comparable speed to a typical sequence similarity search and thus SHOOT is provided as a phylogenetically informative alternative to BLAST, and as a general-purpose sequence search algorithm for analysis and retrieval of related biological sequences.

Local similarity or profile-based search methods such as BLAST [11], DIAMOND [5] or MMseqs [18] have a wide range of uses across the biological and biomedical sciences. The near-ubiquitous utility of these methods has led to them being referred to as the Google of biological research. However, one of the most frequent use cases of these searches is to identify orthologs of a given query sequence. Due to the frequent occurrence of gene duplication and loss, orthologs are often indistinguishable from paralogs in the results of local similarity searches. This is because a given query sequence can have none, one, or many orthologs in a related species. Accordingly, the sequences identified by local similarity searching methods will be an unknown mixture of orthologs and paralogs [19]. The problem of distinguishing orthologs from paralogs can be partially mitigated by a reciprocal best hit search, but with low recall [19]. Phylogenetic methods are required to correctly distinguish orthologs from paralogs as they are readily able to distinguish sequence similarity (branch length) and evolutionary relationships (the topology of the tree).

SHOOT was designed to provide the accuracy and information of a phylogenetic analysis with the speed and simplicity of a local sequence similarity search. By pre-computing the within-database sequence relationships, SHOOT can perform an individual search in a comparable time to BLAST. However, instead of a returning a list of similar sequences SHOOT provides a full maximum-likelihood phylogenetic tree as a result enabling immediate phylogenetic interrogation of the sequence search results. A phylogenetic tree provides the best representation available of the evolutionary history of a gene family. The tree allows the identification of speciation and gene duplication events and thus the identification of orthologs and paralogs. While, SHOOT identifies orthologs and paralogs algorithmically the phylogenetic tree can and should also be examined by a user to gain an understanding of how the gene family has evolved, using the orthology assignment by SHOOT as a guide.

A standard phylogenetic approach to identifying orthologs of a query gene is to begin a local sequence similarity search or profile search (HMMER [20], MMseqs [18]). Frequently, an e-value cut-off is applied to identify a set of similar sequences for subsequent phylogenetic analysis. Because e-values (and their constituent bit-scores) are imperfectly correlated with evolutionary relatedness, the set of similar sequences meeting the search threshold will often be missing some genes as well as often including genes that should not be present. A systematic study using HMMER found that for all n genes from an orthogroup clade to pass an e-value threshold, on average the threshold would have to be set such that 1.8n genes in total met the threshold [21]. i.e. an additional 80% of genes needed to be included, on average, to ensure the orthogroup was complete [21]. Thus, unless a very lenient search is used, genes will be incorrectly absent from the final tree. This can lead to incorrect rooting and subsequent mis-interpretation even by phylogenetic experts [21]. Thus, even for bespoke phylogenetic analyses, it is better to use phylogenetic methods to first select the clade of genes of interest. SHOOT supports this by inferring the tree for the entire family of detectable homologs. The use of trees for complete sets of homologs, together with the use of OrthoFinder’s robust tree-rooting algorithm [13], avoids the problem of mis-rooting and misinterpretation of a tree inferred for a more limited set of genes. Also, by using OrthoFinder clustering approach [13, 22], hits missed for a single sequence are also corrected by multiple hits identified for its homologs. This “phylogenetic gene selection workflow” is supported by SHOOT’s web interface, which allows a clade of genes to be selected and the protein sequences for just this clade to be downloaded for downstream user analyses.

In summary, SHOOT was designed to be as easy to use as BLAST, but to provide phylogenetically resolved results in which the query sequence is correctly placed in a phylogenetic tree. In this way the phylogenetic history of the query sequence and its orthologs can be immediately visualised, interpreted, and retrieved.

## Materials and Methods

### Database preparation

SHOOT consists of a database preparation program and a database search program. The database preparation program takes as input the results of an OrthoFinder [13] analysis of a set of proteomes.

To prepare phylogenetic databases for the SHOOT website, the OrthoFinder version 3.0 option, “-c1”, was used to cluster genes into groups consisting of all homologs, rather than the default behaviour which is to split homologous groups at the level of orthogroups. The advantage of the creating complete homologous groups is that their gene trees show an expanded evolutionary history of those genes, including ancient gene duplication events linking gene families, rather than only reaching back to the last common ancestor of the included species. This differs from a default OrthoFinder orthogroup analysis, for which the partitioning of genes into taxonomically comparable orthogroups groups is the priority. OrthoFinder-inferred rooted gene trees for these homolog groups are computed using MAFFT [23] and IQ-TREE [24] by using the additional options “-M msa -A mafft -T iqtree -s species_tree.nwk”, where “species_tree.nwk” was the rooted species tree for the included species. For IQ-TREE, the best fitting evolutionary model was tested for using “-m TEST” and bootstrap replicates performed using “-bb 1000”.

The OrthoFinder results were converted to a SHOOT database in two steps: splitting of large trees and creation of the DIAMOND profiles database for assigning novel sequences to their correct gene tree. Large trees are split since the time requirements for adding a sequence to an MSA for a homologous group and for adding a sequence to its tree can grow super-linearly in the size of the group, leading to needlessly long runtimes. It was found that DIAMOND could instead be used to assign a gene to its correct subtree and then phylogenetic placement could be applied to assign the gene to its correct position within the subtree (Figure 4).

The script “split_large_tree.py” was used to split any tree larger than 2500 genes into subtrees of no more than 2500 genes each. Each subtree tree also contained an outgroup gene, from outside the clade in the tree for that subtree, which was required for the later sequence search stage. For each tree that was split into subtrees, a super-tree was also created by the script of the phylogenetic relationships linking the subtrees. For each subtree, the script extracted the sub-MSA for later use. This subtree size of 2500 genes was chosen as it is the approximate upper limit tree size for which SHOOT could place a novel query gene in the tree in 15 seconds. This was judged to be a reasonable wait for users of the website to receive the tree for their query sequence. For the databases provided by the SHOOT website, between 2 and 40 of the largest trees were split into subtrees.

The script “create_shoot_db.py” was used to create a DIAMOND database of “profiles” for each unsplit tree or each subtree. A profile here refers to a set of representative sequences that best describe the sequence variability within a homologous group. These profiles are used to assign a novel query sequence to the correct tree or subtree. The representative sequences for a gene tree are selected using k-means clustering applied to the MSA corresponding to that (sub)tree using the python library Scikit-learn [25]. For each cluster, the sequence closest to the centroid is chosen as a representative. For a homologous group of size N genes, k=N/10 representative sequences are used, with a minimum of min(20, N) representative sequences. This ensures that large and diverse homologous groups have sufficient representative sequences in the assignment database.

### Database search

A query sequence is searched against the profiles database using DIAMOND [5] with default sensitivity and an e-value cut-off of 10^-3^. If no hit is found, a second search is performed with the “--ultra-sensitive” setting. The top hitting sequence is used to assign the gene to the correct tree or subtree. The query gene is added to the pre-computed alignment using the MAFFT “--add” option and a phylogenetic tree is computed from this alignment using the precomputed tree for the reference alignment using EPA-ng [12] and gappa [26].

If the gene is added to a subtree then the tree is rooted on the outgroup sequence for that subtree. The outgroup is then removed from the subtree and the subtree is grafted back into the original larger tree, using the supertree to determine the overall topology. This method provides the accuracy of phylogenetic analysis to place the gene in its correct position within the subtree while at the same time providing the user with the full gene history for the complete homologous group given by the supertree, which was calculated in full in the earlier database construction phase. All tree manipulations by SHOOT are performed using the ETE Toolkit [27].

### Curated databases

For the Plants database, the protein sequences derived from primary transcripts were downloaded from Phytozome [28]. The Uniport Reference Proteomes database was constructed using the 2020 Reference Proteomes [17]. For the Fungi and Metazoa databases the proteomes were downloaded from Ensembl [29] and the longest transcript variant of each gene was selected as a representative of that gene using OrthoFinder’s “primary_transcripts.py” script [13]. The Bacterial and Archaeal database proteomes were downloaded from UniProt [30]. The parallelisation of tasks in the preparation of the databases was performed using GNU parallel [31].

### Accuracy validation & performance

The UniProt Reference Proteomes database was used for validation of the SHOOT phylogenetic placements using a leave-one-out test. As this database covers the greatest phylogenetic range (covering all domains of life), its homologous groups contain the greatest sequence variability, and it provides the severest test of the accuracy of SHOOT. Test cases were constructed by selecting 1000 ‘cherries’ (pairs of genes sister to one another) with 95% bootstrap support from gene trees with median bootstrap support of at least 95%. The use of cherries allowed BLAST to be tested alongside SHOOT. This test was possible for BLAST since it would only have to identify a single closest gene, rather than having to identify a gene as the sister gene to a whole clade of genes (as SHOOT is designed to be able to do). The bootstrap support criteria ensured that the correct result was known with high confidence so that both methods could be assessed accurately. To ensure an even sampling of test cases, at most one test case was extracted from any one gene tree. Both the BLAST and SHOOT databases were completely pruned of the 1000 test cases. Each of the 1000 test cases was run using 16 cores of an Intel Xeon E5-2683 CPU and the runtime recorded (Figure 2).

To calculate the Mean Average Precision at k score, the expected trees were re-inferred using RAxML with the best-fitting model [32] so that a different method were used to that used in the SHOOT database construction. For each test gene the ordered list of closest homologs was calculated using branch length distance in the SHOOT results trees and e-values (with ties broken by bit score) for the BLAST results. These ordered homologs were compared to the expected ordered list of closest homologs from the expected RAxML trees to calculate the precision at each value of k from 1 to 50 and these precision scores were averaged over the 1000 test cases.

The ortholog prediction accuracy tests calculated the precision, recall and F-score for identifying orthologs in *Homo sapiens* for genes from *Mus musculus, Gallus gallus, Danio rerio, Ciona intestinalis, Drosophila melanogaster* and *Saccharomyces cerevisiae*. For each of these 6 species 100 genes were sampled at random. The expected orthologs were obtained from OrthoFinder 2020 Quest for Orthologs benchmark results, obtained from the benchmarking server: https://orthology.benchmarkservice.org. For SHOOT, the orthologs were inferred using the species-overlap method [15] on the SHOOT results trees. For BLAST orthologs were predicted using the best hit (BH) method and the reciprocal best hit (RBH) method using the e-value scores.

### SHOOT website

The tree visualisation is provided by the phylotree.js library [33]. The SHOOT website is implemented in JavaScript and Bootstrap and using the Flask web framework.

## Declarations

### Ethics approval and consent to participate

Not applicable

### Consent for publication

Not applicable

### Availability of data and material

The SHOOT source code is available at https://github.com/davidemms/SHOOT. The code for the SHOOT webserver is available at https://github.com/davidemms/SHOOT_webserver. A compressed archive of all data is available at the Zenodo research data archive at https://doi.org/10.5281/zenodo.5602736 [34]. A webserver running SHOOT is available at https://shoot.bio.

### Authors’ contributions

DE and SK conceived and designed the project. DE developed the algorithms. DE and SK discussed the results and wrote the manuscript. All authors read and approved the final manuscript.

## Acknowledgements

The authors would like to thank the members of the Department of Plant Sciences at the University of Oxford and the SHOOT user community for their feedback on the initial versions of the method and webserver.

---

**Supplementary Table 1:**
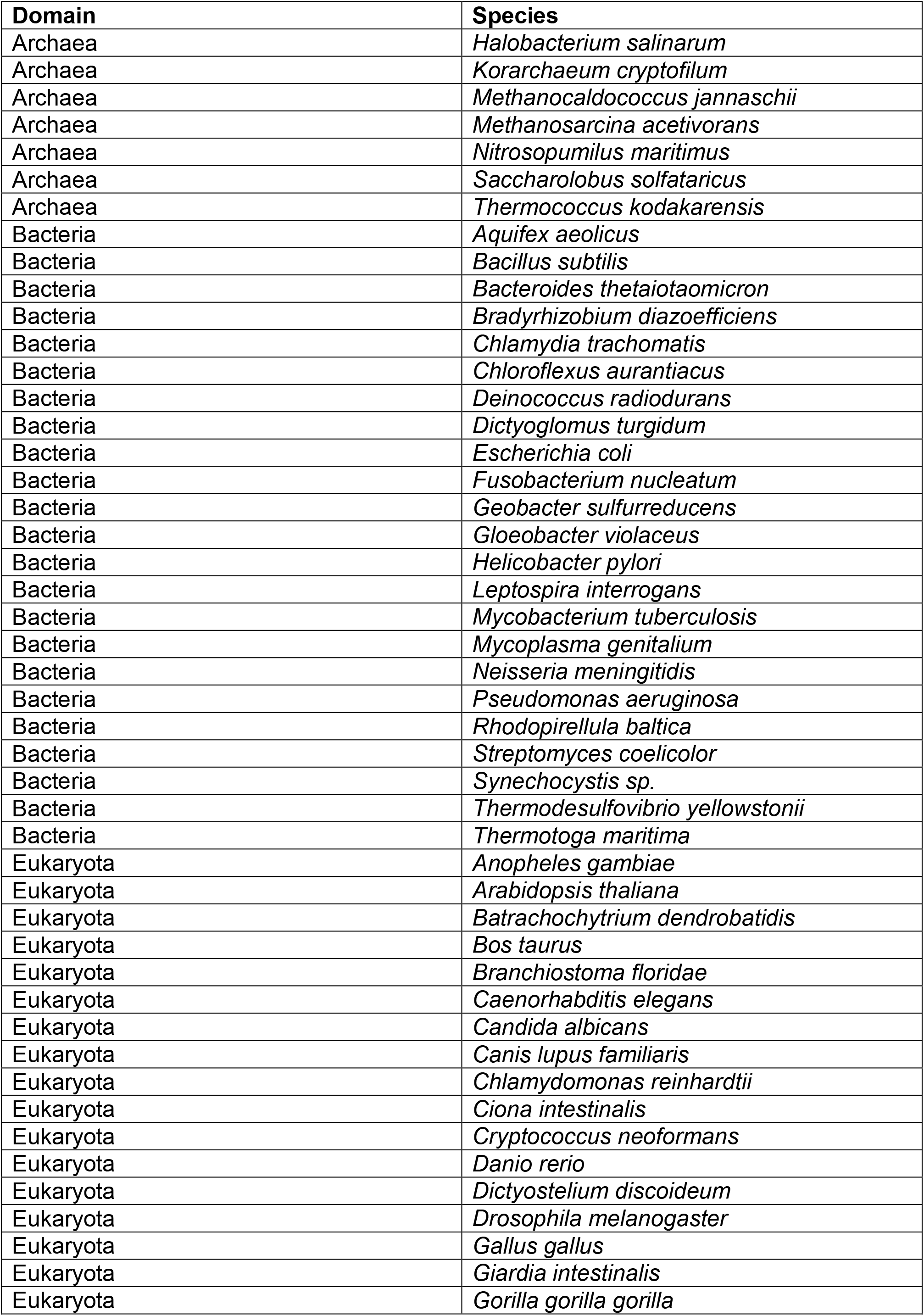
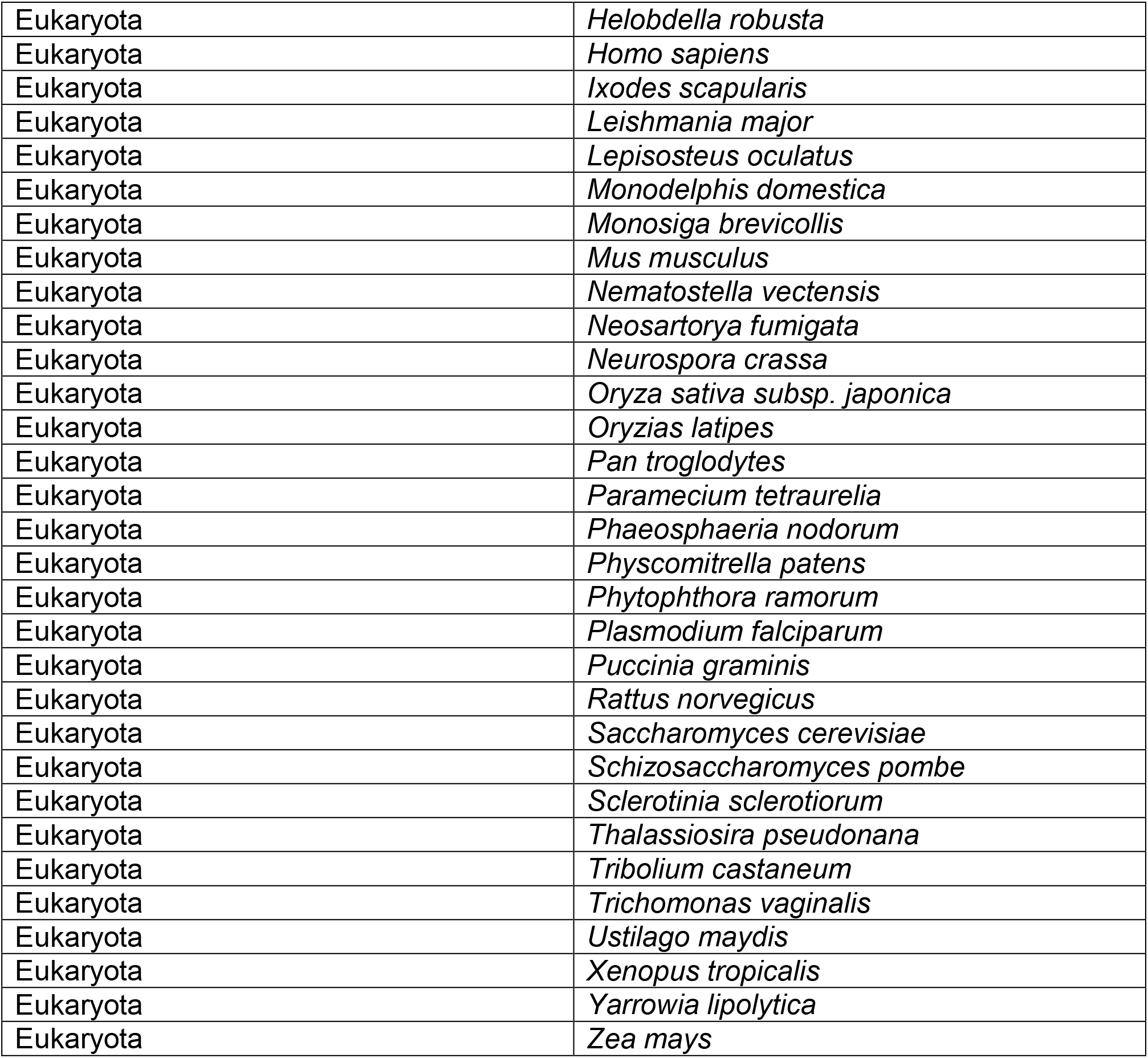
UniProt 2020 Reference Proteomes – Species list.

**Supplementary Table 2:**
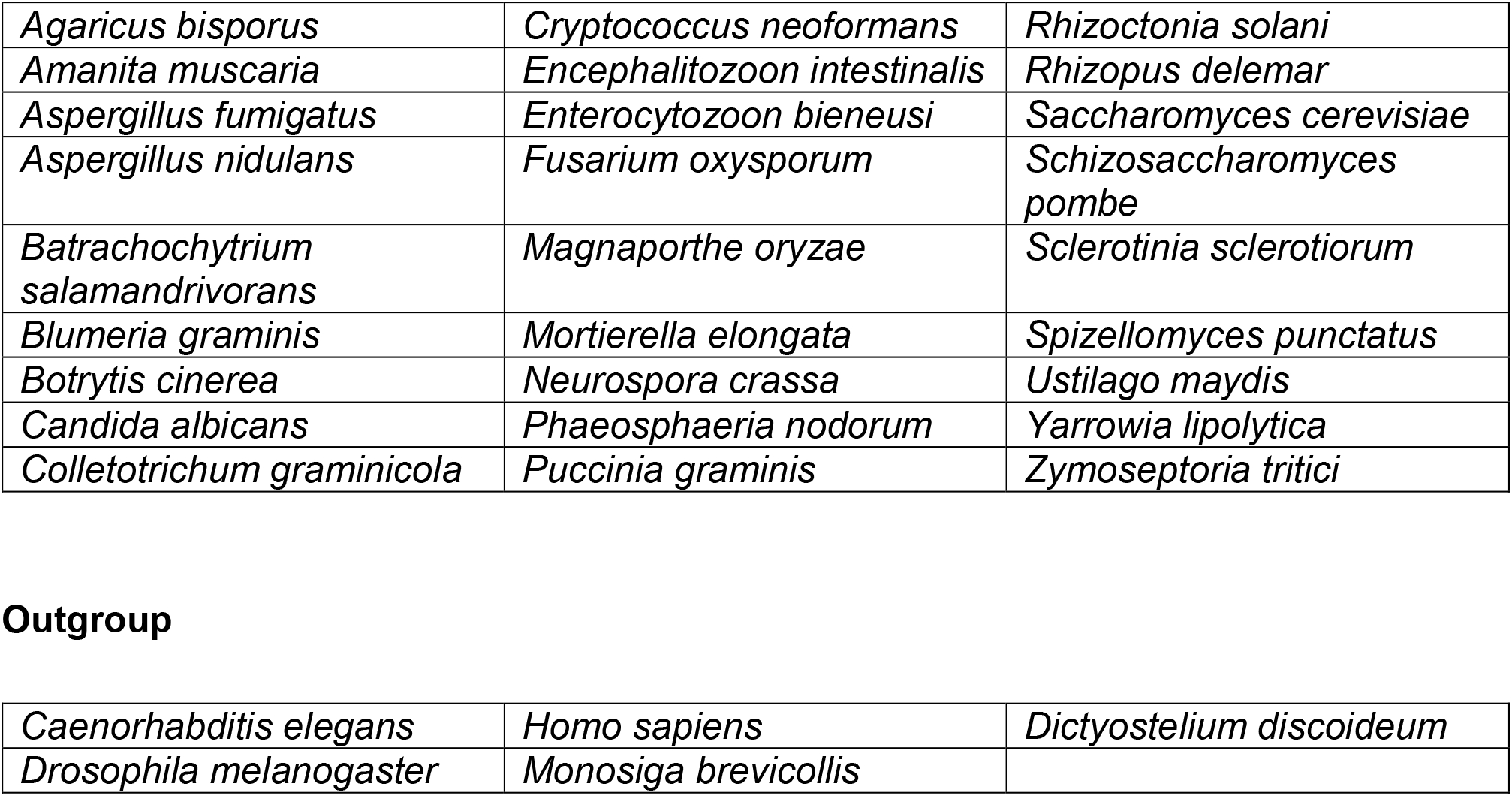
Fungi species list.

|  |  |  |
| --- | --- | --- |
| *Agaricus bisporus* | *Cryptococcus neoformans* | *Rhizoctonia solani* |
| *Amanita muscaria* | *Encephalitozoon intestinalis* | *Rhizopus delemar* |
| *Aspergillus fumigatus* | *Enterocytozoon bieneusi* | *Saccharomyces cerevisiae* |
| *Aspergillus nidulans* | *Fusarium oxysporum* | *Schizosaccharomyces pombe* |
| *Batrachochytrium salamandrivorans* | *Magnaporthe oryzae* | *Sclerotinia sclerotiorum* |
| *Blumeria graminis* | *Mortierella elongata* | *Spizellomyces punctatus* |
| *Botrytis cinerea* | *Neurospora crassa* | *Ustilago maydis* |
| *Candida albicans* | *Phaeosphaeria nodorum* | *Yarrowia lipolytica* |
| *Colletotrichum graminicola* | *Puccinia graminis* | *Zymoseptoria tritici* |
| **Outgroup** |  |  |
| *Caenorhabditis elegans* | *Homo sapiens* | *Dictyostelium discoideum* |
| *Drosophila melanogaster* | *Monosiga brevicollis* |  |

**Supplementary Table 3:**
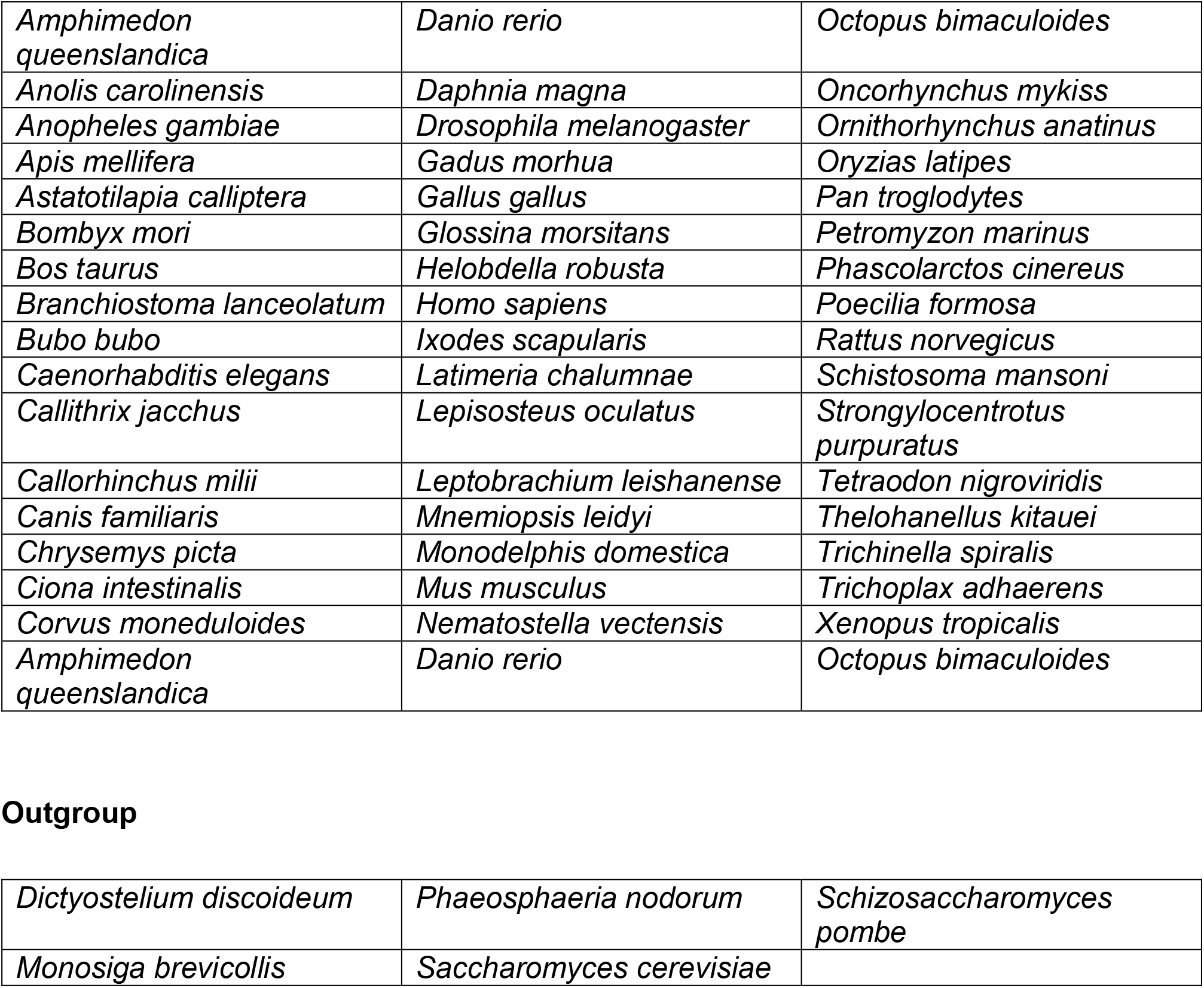
Metazoan species list.

|  |  |  |
| --- | --- | --- |
| *Amphimedon queenslandica* | *Danio rerio* | *Octopus bimaculoides* |
| *Anolis carolinensis* | *Daphnia magna* | *Oncorhynchus mykiss* |
| *Anopheles gambiae* | *Drosophila melanogaster* | *Ornithorhynchus anatinus* |
| *Apis mellifera* | *Gadus morhua* | *Oryzias latipes* |
| *Astatotilapia calliptera* | *Gallus gallus* | *Pan troglodytes* |
| *Bombyx mori* | *Glossina morsitans* | *Petromyzon marinus* |
| *Bos taurus* | *Helobdella robusta* | *Phascolarctos cinereus* |
| *Branchiostoma lanceolatum* | *Homo sapiens* | *Poecilia formosa* |
| *Bubo bubo* | *Ixodes scapularis* | *Rattus norvegicus* |
| *Caenorhabditis elegans* | *Latimeria chalumnae* | *Schistosoma mansoni* |
| *Callithrix jacchus* | *Lepisosteus oculatus* | *Strongylocentrotus purpuratus* |
| *Callorhinchus milii* | *Leptobranchium leishanense* | *Tetraodon nigroviridis* |
| *Canis familiaris* | *Mnemiopsis leidyi* | *Thelohanellus kitauei* |
| *Chrysemys picta* | *Monodelphis domestica* | *Trichinella spiralis* |
| *Ciona intestinalis* | *Mus musculus* | *Trichoplax adhaerens* |
| *Corvus moneduloides* | *Nematostella vectensis* | *Xenopus tropicalis* |
| *Amphimedon queenslandica* | *Danio rerio* | *Octopus bimaculoides* |
| **Outgroup** |  |  |
| *Dictyostelium discoideum* | *Phaeosphaeria nodorum* | *Schizosaccharomyces pombe* |
| *Monosiga brevicollis* | *Saccharomyces cerevisiae* |  |

**Supplementary Table 4:**
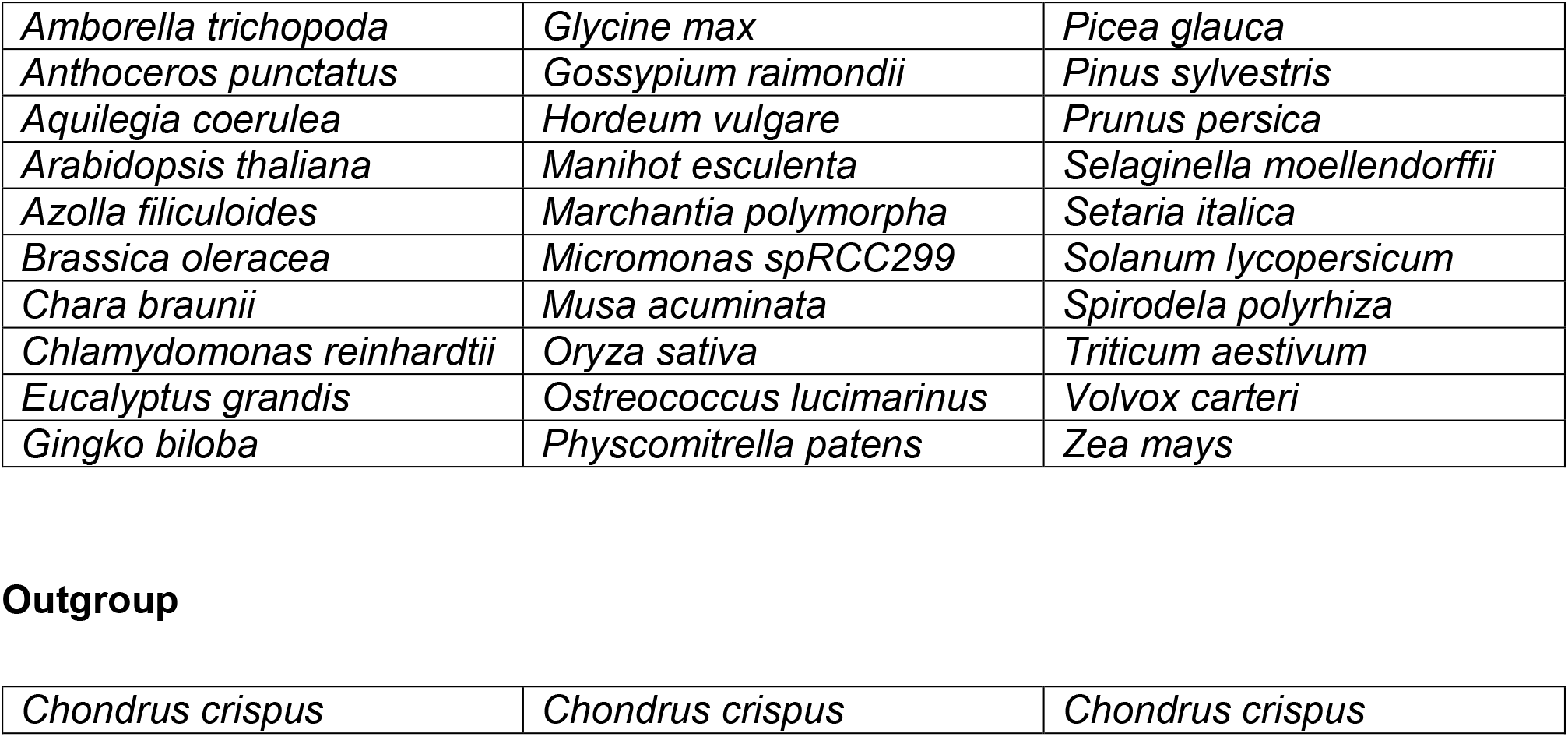
Plants species list.

|  |  |  |
| --- | --- | --- |
| *Amborella trichopoda* | *Glycine max* | *Picea glauca* |
| *Anthoceros punctatus* | *Gossypium raimondii* | *Pinus sylvestris* |
| *Aquilegia coerulea* | *Hordeum vulgare* | *Prunus persica* |
| *Arabidopsis thaliana* | *Manihot esculenta* | *Selaginella moellendorffii* |
| *Azolla filiculoides* | *Marchantia polymorpha* | *Setaria italica* |
| *Brassica oleracea* | *Micromonas spRCC299* | *Solanum lycopersicum* |
| *Chara braunii* | *Musa acuminata* | *Spirodela polyrhiza* |
| *Chlamydomonas reinhardtii* | *Oryza sativa* | *Triticum aestivum* |
| *Eucalyptus grandis* | *Ostreococcus lucimarinus* | *Volvox carteri* |
| *Ginkgo biloba* | *Physcomitrella patens* | *Zea mays* |
| **Outgroup** |  |  |
| *Chondrus crispus* | *Chondrus crispus* | *Chondrus crispus* |

**Supplementary Table 5:**
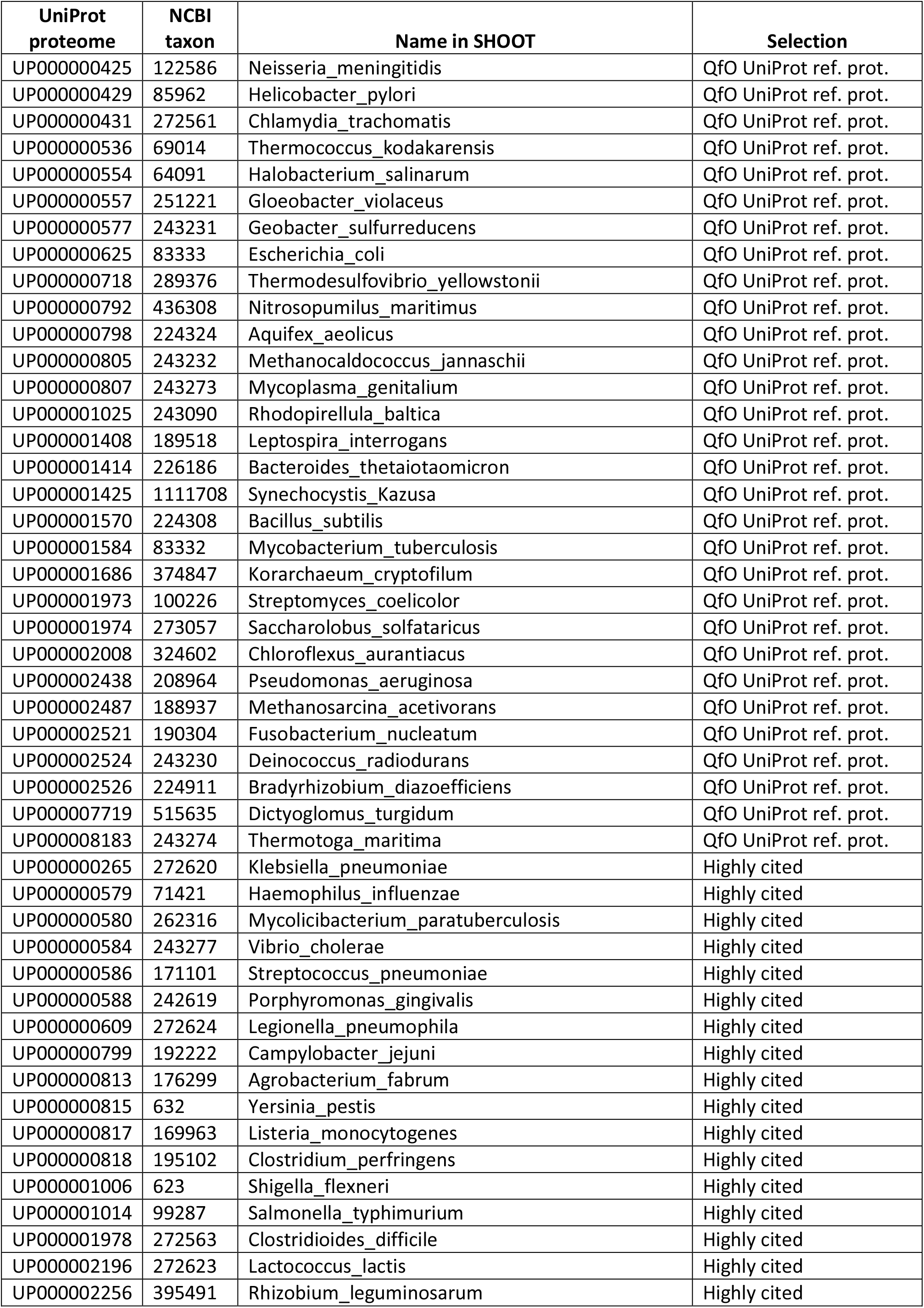
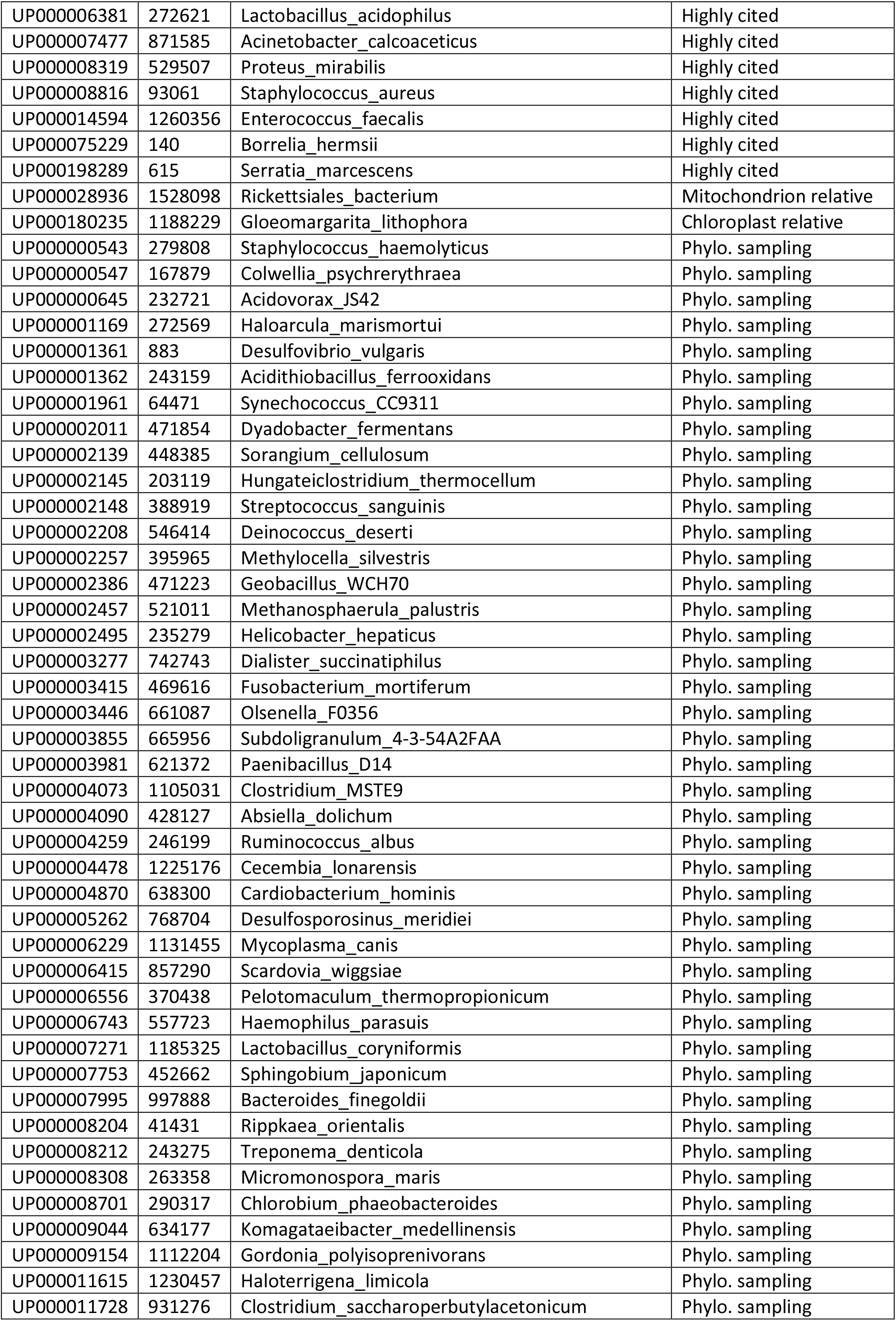
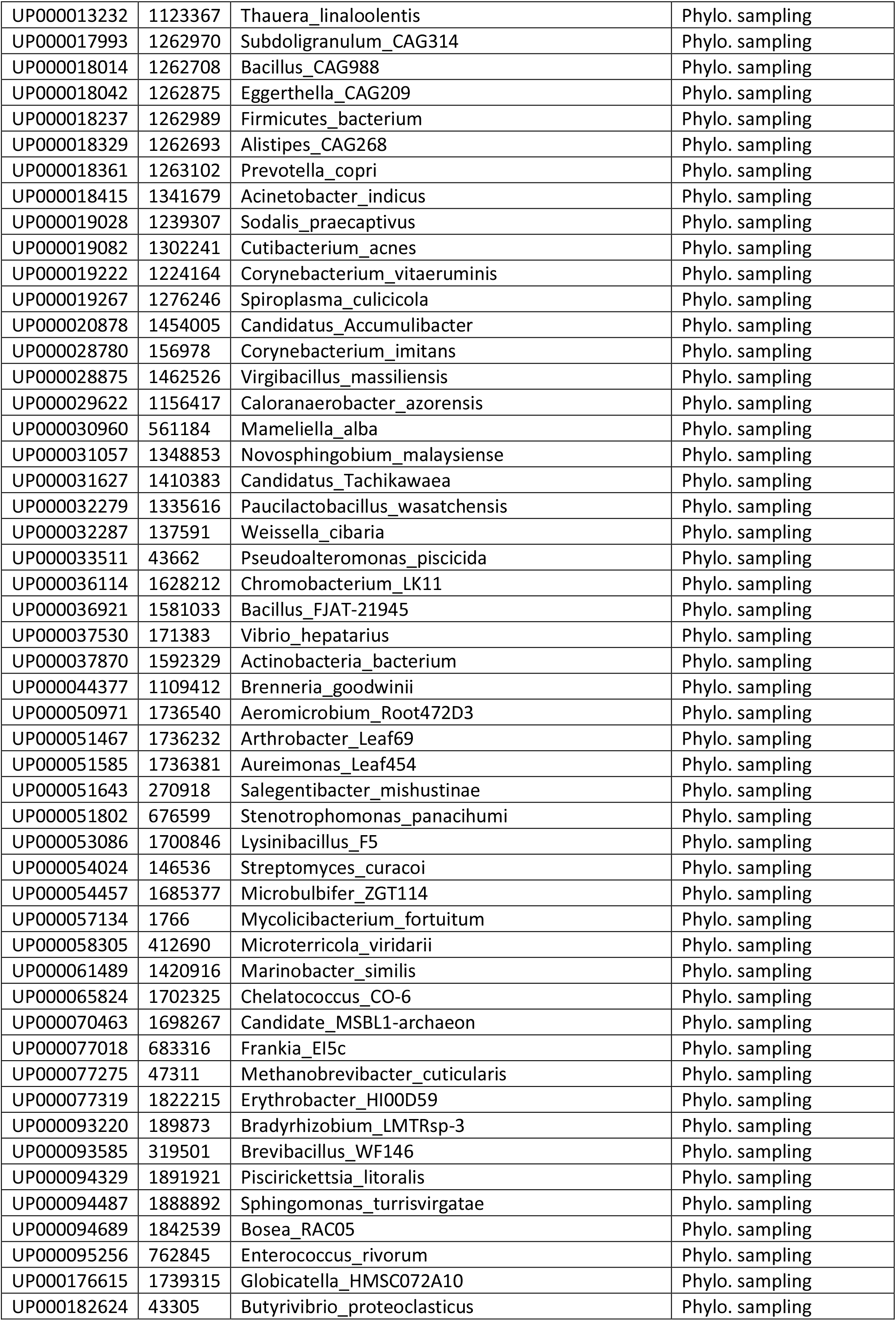
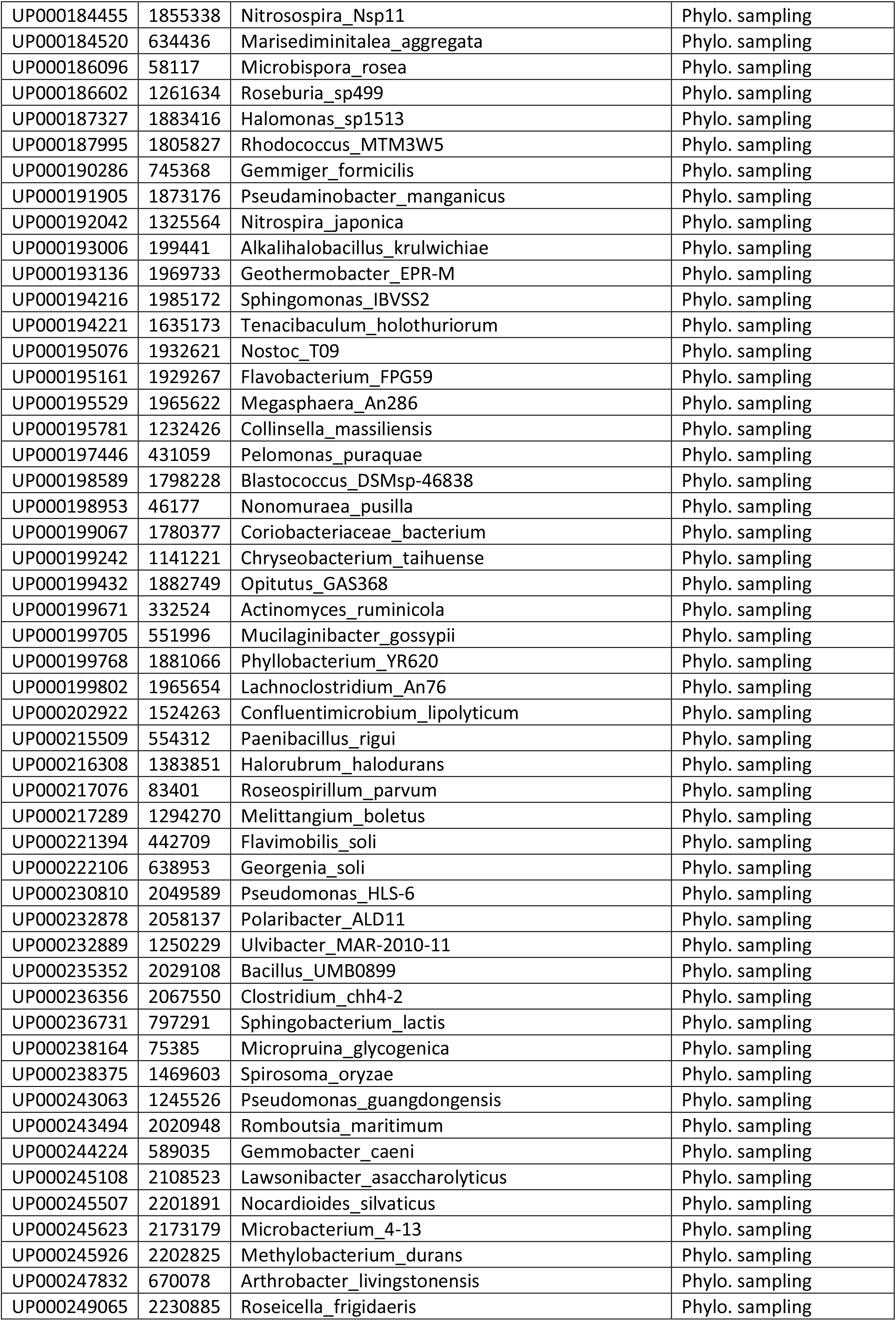
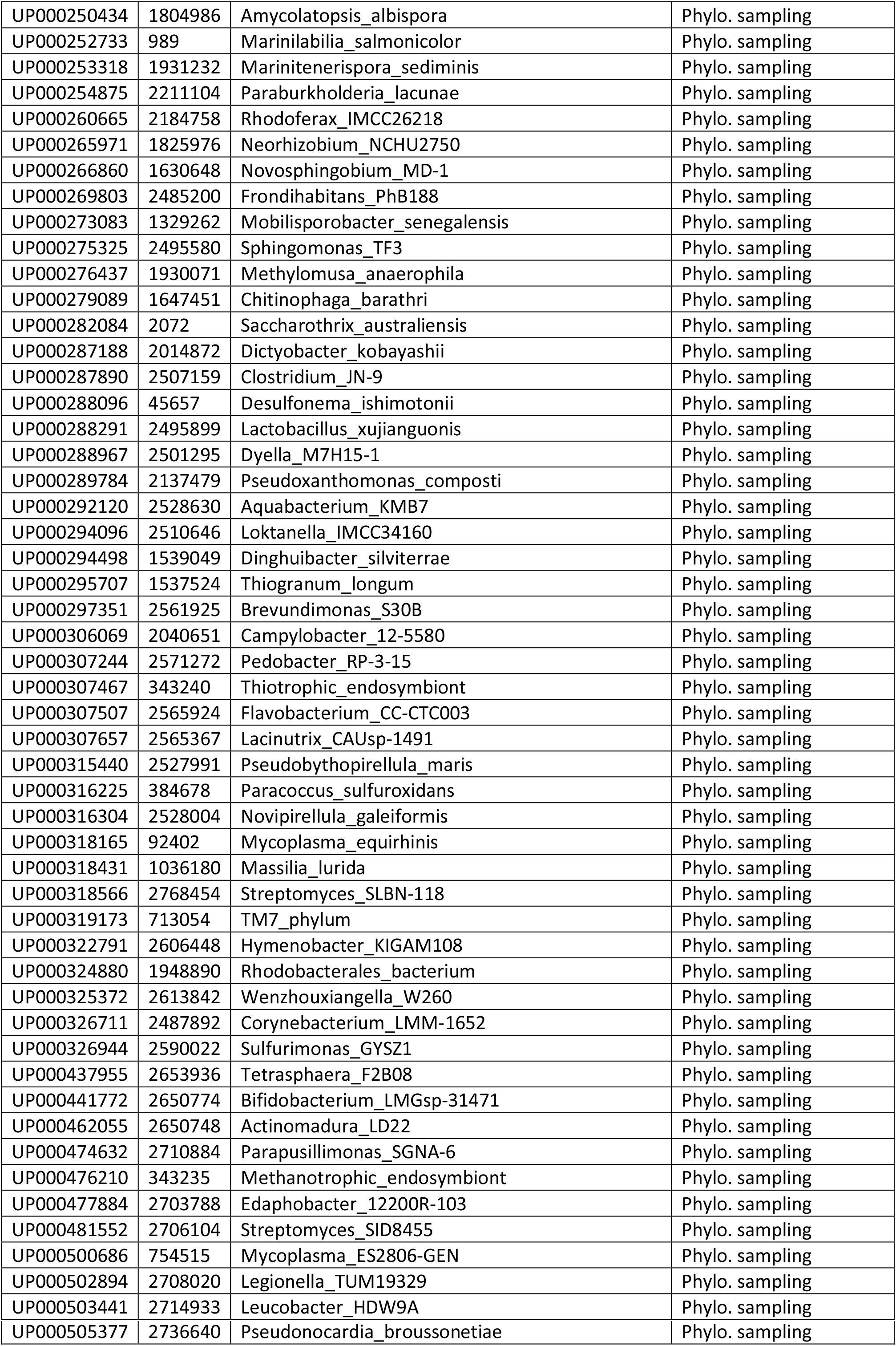
Bacterial & Archaeal strains list.

## Supplementary Figure Legends

**Supplementary Figure 1.**
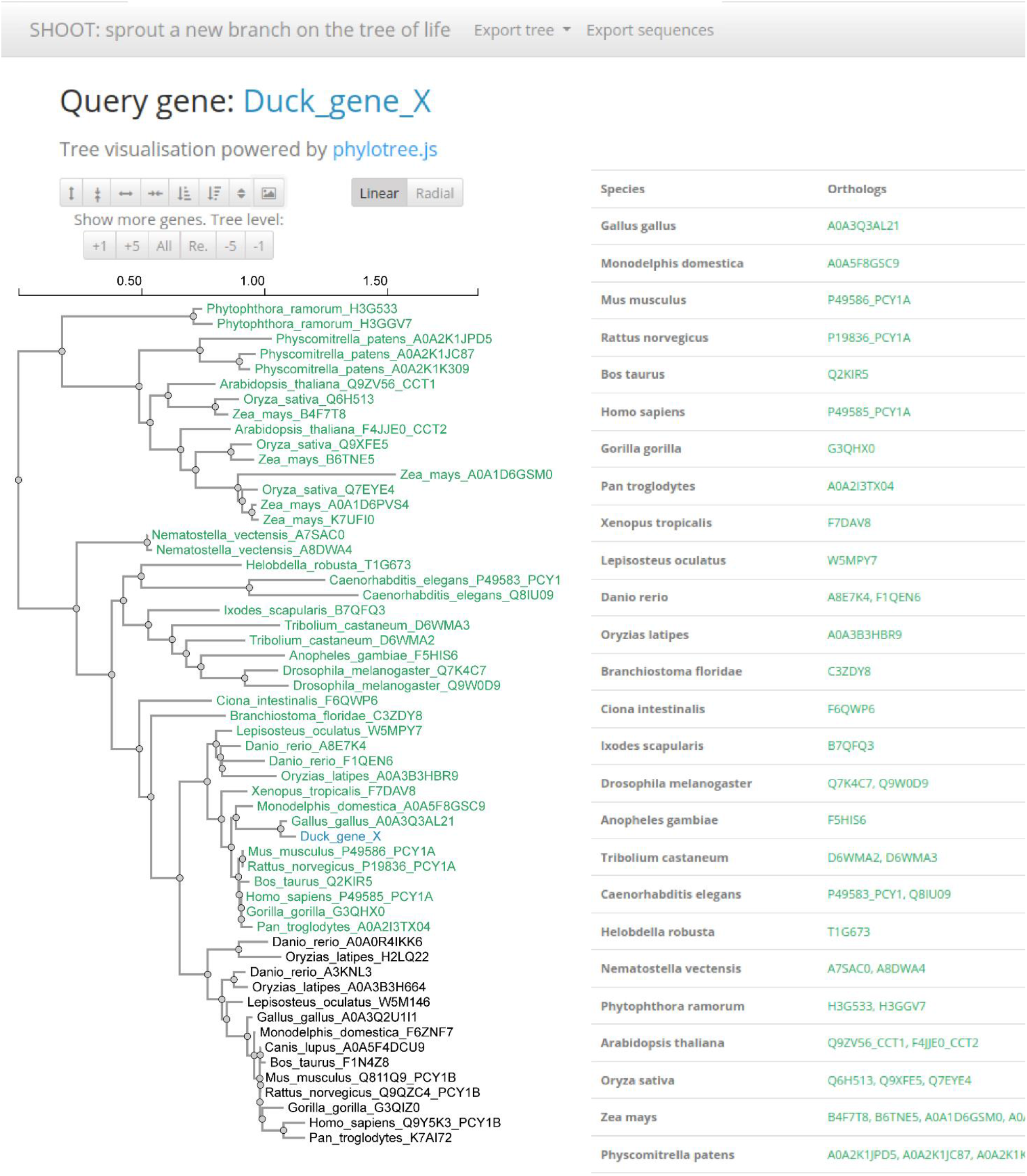
An example gene tree and orthologs table returned by SHOOT. Here, the UniProt Reference Proteomes database was searched using a for a query gene sequence labelled “Duck_gene_X”. This corresponds to the Duck protein ENSAPLP00000002788, which is not included in the database.

**Supplementary Figure 2.**
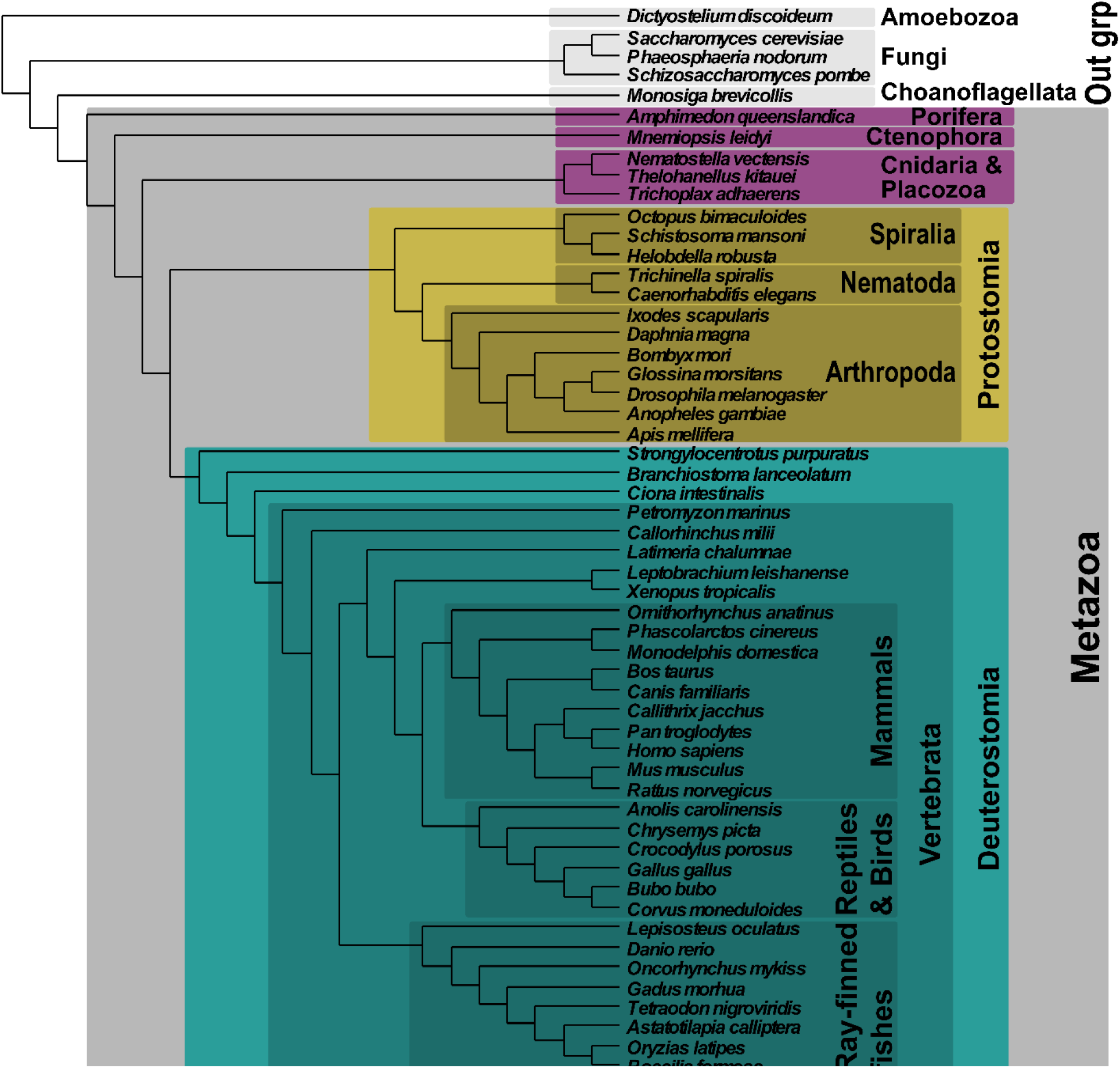
Phylogeny for the species in the Metazoan dataset.

